# Digital phenotyping by consumer wearables identifies sleep-associated markers of cardiovascular disease risk and biological aging

**DOI:** 10.1101/527077

**Authors:** Jing Xian Teo, Sonia Davila, Chengxi Yang, Chee Jian Pua, Jonathan Yap, Swee Yaw Tan, Anders Sahlén, Calvin Woon-Loong Chin, Bin Tean Teh, Steven G Rozen, Stuart Alexander Cook, Khung Keong Yeo, Patrick Tan, Weng Khong Lim

## Abstract

Despite growing adoption of consumer wearables, the potential for sleep metrics from these devices to contribute to sleep-related biomedical research remains largely uncharacterized. Here we analyze sleep tracking data, along with questionnaire responses and multi-modal phenotypic data, generated from 482 normal volunteers. First, we provide a detailed comparison of wearable-derived and self-reported sleep metrics, particularly total sleep time (TST) and sleep efficiency (SE). We then identified demographic, socioeconomic and lifestyle factors associated with wearable-derived sleep duration. We also analyzed our multi-modal phenotypic data and showed that wearable-derived TST and SE are associated with various cardiovascular disease risk markers, whereas self-reported measures were not. Using whole-genome sequencing data, we estimated leukocyte telomere length and showed that volunteers with insufficient sleep also exhibit premature telomere attrition. Our study highlights the potential for sleep metrics generated by consumer wearables to provide novel insights into data generated from population cohort studies.

## Introduction

Relationships between sleep and health outcomes have been described by many studies over the years. Among others, insufficient sleep has been associated with obesity[1,2], hypertension [3–6], cardiovascular disease (CVD) [7–10], insulin resistance [11–14] and even premature death [15,16]. Previous studies on sleep-health interactions have relied on three methods to quantify sleep; sleep questionnaires/diaries, actigraphy and polysomnography (PSG). There are drawbacks associated with each approach: 1) sleep questionnaires/diaries lack precision and rely on subjective recall [17], 2) actigraphy involves specialized devices and are only suitable for relatively short studies, and 3) PSG studies while being the gold-standard in accuracy are very resource-intensive to conduct [18].

The digital revolution has resulted in the proliferation of consumer wearables with activity tracking functionality. Shipments of wearables exceeded 114 million units in 2017 and is expected to double by 2021 [19,20]. Although many of these wearables are marketed either as fitness trackers or smartwatches, the line between these two categories is rapidly blurring. Beyond physical activity, such devices can also track sleep, with more advanced models able to provide sleep staging information via integrated heart rate (HR) sensors. Although consumer wearables are marketed as tools to promote healthy sleep habits, their widespread availability and growing adoption suggest the possibility of using them as a source of quantitative sleep data for sleep and health research.

Biomedical researchers are beginning to explore the potential of sleep data derived from consumer wearables. First, researchers have compared the accuracy of sleep as measured by consumer wearables from several manufacturers (e.g. Fitbit and Jawbone) with gold standard PSG measurements [21–24]. They found that consumer wearables were reasonably sensitive in detecting sleep but performed poorly in detecting periods of wakefulness. Compared to research actigraphs, consumer wearables have similar sensitivity (sleep detection) and specificity (wake detection) [23,25]. Some cohort studies have begun using consumer wearables. For example, we recently used Fitbit-derived sleep tracking data to show differences in sleep patterns among volunteers stratified into various activity pattern clusters [26]. Xu et al. used Fitbit Charge HR sleep data from 748 individuals to demonstrate independent associations between both sleep duration and sleep duration variation with body mass index (BMI) [27]. Additionally, Turel et al. used Fitbit devices to show a negative association between sleep duration and abdominal obesity [28].

Despite these advances, various gaps remain in our understanding of the utility of sleep data from consumer wearables in biomedical research. First, there is little information on how sleep metrics from consumer wearables compare with self-reported sleep quality from questionnaires such as the Pittsburgh Sleep Quality Index (PSQI), which is typically used in large cohort studies where it is it impractical and costly to use actigraphy or PSG to measure individual sleep parameters [27,29]. Such a comparison is important if consumer wearables are to be considered as replacements or adjuncts of sleep questionnaires in future cohort studies. Second, there has yet to be an analysis of consumer wearable sleep data in the context of a comprehensive multi-modal dataset comprising various health parameters such as anthropometry, blood pressure, lipid profile, cardiac imaging, especially in Asians; a population known to have considerably different sleep behavior compared to Western cohorts [27,30].Third, although insufficient sleep has been linked to accelerated aging [31–34], this phenomenon has yet to be explored in a sizeable cohort using consumer wearables. Finally, there has yet to be an exploration of how wearable sleep data correlates with demographic, socioeconomic and lifestyle factors. Using an expanded cohort and dataset compared to our initial study [26], we sought to address these gaps through a comprehensive analysis of sleep data derived from Fitbit Charge HR activity trackers worn by 482 Singaporean volunteers. Apart from the wearable tracking, these volunteers have been comprehensively profiled for CVD risk markers including anthropometrics, blood pressure, lipid panel and fasting blood glucose (FBG).

## Results

### Comparison between wearable-derived and subjective sleep metrics

The cohort of 482 volunteers underwent a wearable study lasting an average of 4 days, where Fitbit Charge HR activity trackers were used to track physical activity, HR and sleep. Summary statistics for the cohort are shown in **Table 1** (full details in **S1 Data**). At time of enrollment, the volunteers were on average 46 years of age (range 21 – 69 years). These volunteers on average had 4 nights of tracked sleep (range 3 – 11 nights), with a mean total sleep time (TST) of 6 hours and 28 minutes. The first analysis that we performed was to compare objective sleep measures obtained using consumer wearables to subjective sleep measures in the form of PSQI responses. The PSQI questionnaire comprises various sections, each containing questions that seek to quantify a different aspect of sleep quality (e.g. sleep duration, sleep efficiency, sleep latency, subjective sleep quality, etc.). It then summarizes the scores of individual components into a global PSQI score representing overall sleep quality. We first compared wearable-derived TST and SE with global PSQI scores and found correlations in neither (r_s_ = -0.089, p = 0.091 and r_s_ = -0.080 p = 0.129 respectively). Wearable-derived TST, however, showed a significant, albeit weak correlation with self-reported TST (r_s_ = 0.283, p = 2.394×10^-10^). We asked if this weak correlation is at least in part due to the relatively short study duration. We thus modified the inclusion criteria from at least three nights of tracked sleep to four and five nights. Indeed, when the thresholds were increased, correlation with self-reported TST rose correspondingly to 0.322 (p = 6.218×10^-09^, n = 310) and 0.397 (p = 1.425×10^-06^, n = 138) respectively, despite remaining relatively weak. Apart from raw self-reported TST, we also categorized self-reported TST by levels specified in component 3 of the PSQI, which profiles habitual sleep duration. Compared to those with the lowest component 3 score of 0 (> 7 hours of sleep), those with scores of 1 (6 -7 hours) and 2 (5-6 hours) had lower wearable-derived TST (β = -0.321, p = 0.001 and β = -0.721, p = 4.94×10^-06^ respectively, **Fig 1A**). However, those with a score of 3 (< 5 hours) exhibited no significant difference in TST compared to those with a score of 0 (β = -0.428, p = 0.123), indicating lower concordance with wearable-derived TST among those with self-perceived chronic sleep deprivation. This may be due to the limited number of volunteers in that category (n = 13) and correspondingly higher variability in wearable-derived TST. Overall, we found that volunteers on average over-estimated their habitual sleep duration by 14 minutes compared to the more objective and quantitative wearable-derived measurements.

**Table 1.** Summary statistics of study volunteers. Test p-values for between gender comparisons are shown: For continuous variables, Student t-test was used, whereas categorical values were evaluated using the chi-squared test. The full dataset is available in S1 Data, and this table was generated by code in S2 Data. Abbreviations: BMI = body mass index; WC = waist circumference; WHtR = waist-to-height ration; BFP = body fat percentage; SMP = skeletal muscle percentage; SBP = systolic blood pressure; DBP = diastolic blood pressure; LDL = low-density lipoprotein; HDL = high-density lipoprotein; TG = triglycerides; LTL = leucocyte telomere length; TST = total sleep time; SE = sleep efficiency; GPPAQ = General Practice Physical Activity Questionnaire; PSQI = Pittsburgh Sleep Quality Index.

| Characteristic | Female (n=262; 54.36%) | Male (n=220; 45.64%) | Test |
| --- | --- | --- | --- |
| Age, years | 46.21 (11.35) | 45.80 (12.70) | 0.703 |
| Ethnicity |  |  | 0.001 |
| Chinese | 247 (94.3) | 192 (87.3) |  |
| Indian | 6 (2.3) | 11 (5.0) |  |
| Malay | 2 (0.8) | 14 (6.4) |  |
| Others | 7 (2.7) | 3 (1.4) |  |
| BMI, kg/m<sup>2</sup> | 22.83 (3.80) | 24.27 (3.06) | <0.001 |
| WC, cm | 79.01 (11.08) | 86.82 (9.36) | <0.001 |
| WHtR | 0.50 (0.07) | 0.51 (0.05) | 0.073 |
| BFP, % | 33.16 (7.65) | 23.64 (6.48) | <0.001 |
| SMP, % | 35.49 (4.40) | 42.57 (4.25) | <0.001 |
| SBP, mmHg | 122.54 (17.53) | 132.06 (15.04) | <0.001 |
| DBP, mmHg | 73.82 (12.85) | 82.40 (10.88) | <0.001 |
| Total Cholesterol, mmol/l | 5.38 (0.93) | 5.39 (0.97) | 0.922 |
| LDL, mmol/l | 3.34 (0.81) | 3.45 (0.91) | 0.183 |
| HDL, mmol/l | 1.58 (0.32) | 1.36 (0.32) | <0.001 |
| TGs, mmol/l | 1.02 (0.54) | 1.33 (0.80) | <0.001 |
| Glucose, mmol/l | 5.18 (0.50) | 5.39 (0.73) | <0.001 |
| RestingHR, (Fitbit, bpm) | 69.79 (6.37) | 68.23 (6.48) | 0.008 |
| DailySteps, (Fitbit, x1000) | 10349.55 (3466.19) | 11061.09 (3818.57) | 0.033 |
| LTL, bp | -47.72 (443.41) | -57.12 (536.72) | 0.899 |
| GPPAQ Score | 1.31 (1.12) | 1.95 (1.11) | <0.001 |
| Wearable-derived TST, hr | 6.60 (1.00) | 6.32 (0.98) | 0.002 |
| Self-report TST (PSQI Sleep Hour), hr | 6.59 (1.04) | 6.56 (1.00) | 0.796 |
| Wearable-derived SE, % | 93.08 (2.84) | 92.00 (3.22) | <0.001 |
| Self-reported SE (PSQI Component 4 Score), % | 0.26 (0.63) | 0.22 (0.56) | 0.407 |
| Wearable-derived nocturnal awakenings | 2.00 (1.38) | 1.95 (1.57) | 0.755 |
| Self-reported nocturnal awakenings (PSQI Question 5b) | 1.02 (1.06) | 1.13 (1.13) | 0.263 |
| Global PSQI Score | 3.73 (2.36) | 3.78 (2.14) | 0.854 |

**Fig 1.**
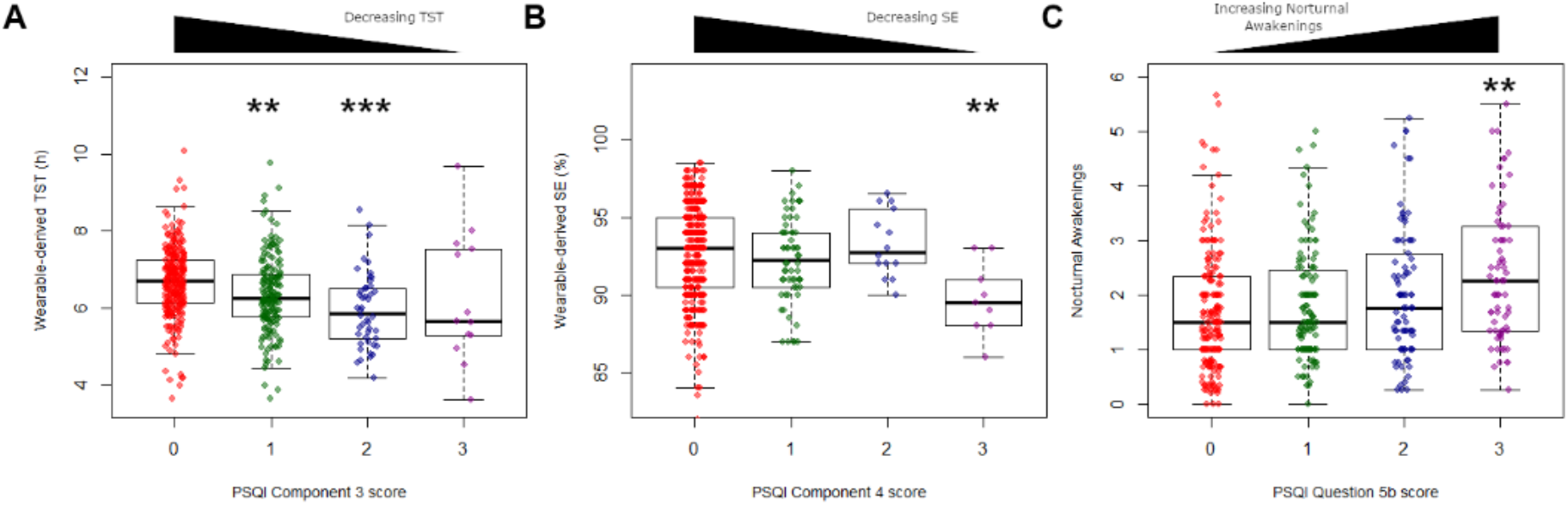
Comparison between wearable-derived and self-reported sleep metrics. (A) Wearable-derived TST and PSQI Component 3 score (sleep duration). (B) Wearable-derived SE and PSQI Component 4 score (sleep efficiency). (C) Wearable-derived nocturnal awakenings and PSQI Component 5b score (nocturnal awakenings). The code to generate this figure can be found in S2 Data. Asterisks denote significance of component score in linear model compared to reference score of 0. ** = p < 0.01, *** = p < 0.001. Abbreviations: TST, total sleep time; SE, sleep efficiency; PSQI, Pittsburgh Sleep Quality Index.

Apart from TST, the Fitbit wearables used in this study also provide sleep efficiency (SE) metrics, which represent fraction of TST over total time in bed. Similarly, the PSQI also attempts to estimate SE based on self-reported habitual sleep duration, sleep times and wake times. An overall comparison between wearable-derived and self-reported SE revealed no correlation (r_s_ = -0.080, p = 0.081). However, when self-reported SE was grouped by PSQI component 4 scoring thresholds (> 85%, 75 - 84%, 65 - 75%, <65%), volunteers in the < 65% group had significantly lower wearable-derived SE compared others (mean SE = 89.722% vs 92.638%, Student’s t-test p-value = 0.005, **Fig 1B**). Thus, only volunteers with the poorest self-perceived SE have concordantly lower wearable-derived SE values. We note that wearable-derived SE is almost uniformly high (mean SE = 92.584%), with little variation (SE standard deviation = 3.060%), which is likely due to the lower sensitivity of Fitbit wearables in detecting wake states as opposed to sleep states [21].

Another wearable-derived sleep metric available was the number of awakenings per sleep session. For each volunteer, we obtained the average daily number of nocturnal awakenings. We then compared this number with responses to question 5b of the PSQI, which asks volunteers how frequently they had trouble sleeping due to waking up in middle of the night or early morning. There was a weak correlation between these values (r_s_ =0.189, p = 4.114×10^-05^), with volunteers reporting the highest (>= 3 times/week) number of nocturnal awakenings having significantly higher daily wearable-detected nocturnal awakenings compared to volunteer reporting no trouble sleeping in the past month (β = 0.586, p = 0.004, **Fig 1C**). Taken collectively, our findings show that sleep metrics obtained from consumer wearables provide objective sleep metrics that, although associated with to a certain extent, are different from subjective measures of sleep quality.

### Relationship between wearable sleep metrics and cohort demographics

We next sought to determine if wearable-derived sleep metrics could be used to identify sleep-associated demographic and behavioral factors in our cohort. To do so, we first considered volunteer information provided through detailed demographic and socioeconomic questionnaires administered during their recruitment. These included basic demographics such as age, gender and ethnicity, but also socioeconomic and lifestyle factors.

Although the volunteers were predominantly of Chinese ethnicity (n = 439, 91.1%), some were of Malay (n = 16, 3.3%), Indian (n = 17, 3.5%) and other (n = 10, 2.1%) ethnicities. Adjusting for age and gender, we found that Malay volunteers on average slept 41 minutes less than Chinese volunteers (β = -40.818, p = 0.009) whereas Indian volunteers slept for an additional 32 minutes compared to their Chinese counterparts (β = 31.907, p = 0.029). Similarly, after adjusting for gender and ethnicity, we found that TST decreased with age (β = -0.493, p = 0.032, **Fig 2B**). For gender, we found that after adjusting for age and ethnicity, female volunteers on average slept 16 minutes longer than their male counterparts (p = 0.005, **Fig 2C**).

**Fig 2.**
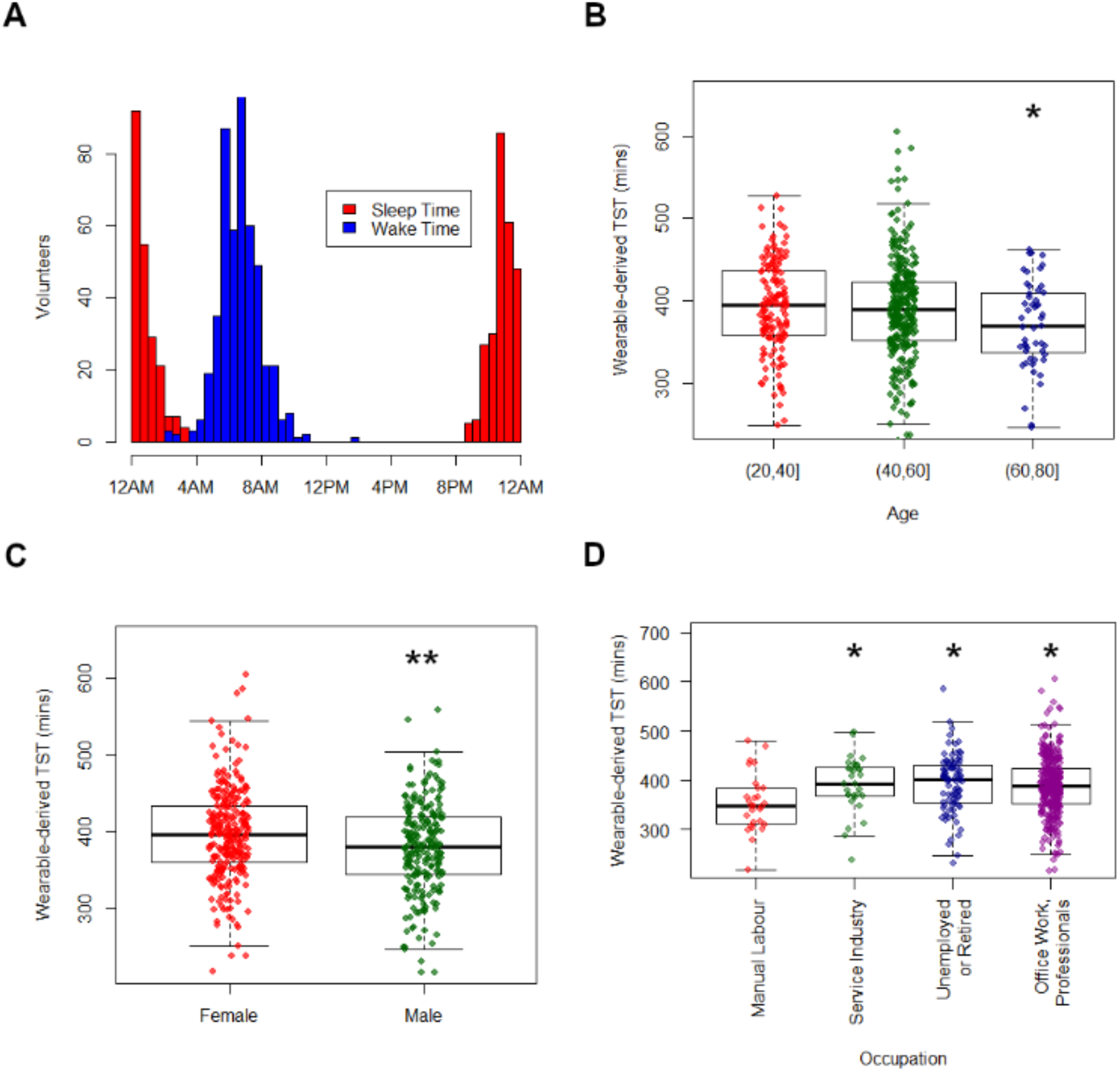
Wearable sleep duration and demographic factors. (A) Distribution of volunteer sleep and wake times. (B-D) Wearable-derived TST by (B) age-group (C) gender (D) occupation type. Asterisks denote significance of factor in linear model compared to reference level (leftmost factor). * = p < 0.05, ** = p < 0.01, *** = p < 0.001. TST = total sleep time.

We next examined relationships between wearable-derived TST and socioeconomic factors. Among others, we considered income levels, residence type, education level and occupation type (**Supplementary Table 1**). Of these factors, occupation type and residence type were associated with TST. Volunteers engaged in manual work slept 27 minutes less then volunteers engaged in other occupation types (i.e. service industry, office work and unemployed/retired, p = 0.022, **Fig 2D**). Furthermore, volunteers living in private residences slept 15 minutes longer than those living in public housing (p = 0.019).

Several self-reported lifestyle factors were also analyzed to determine if they are associated with wearable-derived TST. These included exercise, smoking status, alcohol consumption and caffeine consumption (**Supplementary Table 2**). Apart from alcohol consumption, no other significant associations were found. Volunteers who self-reported alcohol consumption within the past three months slept 19 minutes longer than those who did not, adjusting for age, gender and ethnicity (p =8.54×10^-04^). When alcohol consumption was broken down by type of alcohol, volunteers reporting consumption of hard liquor had the largest difference in TST compared to those that did not (28 minutes longer, p = 0.002), followed by red wine (19 minutes longer, p = 0.008) and beer (18 minutes longer, p = 0.014).

### Associations between wearable-derived sleep metrics and cardiovascular disease markers

A key aim of the study cohort was to study cardiovascular disease (CVD) risk in normal individuals. To that end, various baseline markers of cardiovascular health were collected, including anthropometric measurements (body mass index [BMI], waist circumference [WC], waist-to-height ratio [WHtR]), resting HR, blood pressure (systolic blood pressure [SBP], diastolic blood pressure [DBP]), lipid panel results (total cholesterol [TotalChol], low-density lipoprotein [LDL], high-density lipoprotein [HDL], triglycerides [TG]) and FBG. To determine if wearable-derived sleep metrics were associated with any of these CVD risk markers, we performed multiple linear regression using three different models. Models 1 and 2 consider TST and SE individually, whereas Model 3 includes both TST and SE. All three models included age, gender and daily step counts as covariates. Wearable-derived TST was associated with BMI, TotalChol and resting HR, whereas wearable derived SE was associated with BMI, WC, WHtR and HDL levels (**Table 2**). Neither wearable-derived TST nor SE were significantly associated with blood pressure or FBG levels in this study. We then tested selfreported sleep metrics for TST and SE using the same models and found no significant associations, probably due to the less precise nature of subjective sleep measures. Potential reasons for this are considered in the Discussion.

**Table 2.** Association between wearable-derived sleep metrics and CVD risk markers. Model 1 = TST only, Model 2 = SE only, Model 3 = TST + SE. All models include age and gender as covariates. The code to generate this figure can be found in S2 Data. Highlighted cells are statistically significant (p < 0.05). BMI =body mass index; WC = waist circumference; WHtR = waist-to-height ration; BFP = body fat percentage; SMP = skeletal muscle percentage; SBP = systolic blood pressure; DBP = diastolic blood pressure; TotalChol = total cholesterol; LDL = low-density lipoprotein; HDL = high-density lipoprotein; TG = triglycerides; TST = total sleep time; SE = sleep efficiency. ^a^Marker~Age+Gender+Ethnicity+AverageDailyTotalSteps+Wearable-derived TST ^b^Marker~Age+Gender+Ethnicity+AverageDailyTotalSteps+Wearable-derived SE ^c^Marker~Age+Gender+Ethnicity+AverageDailyTotalSteps+Wearable-derived TST+ Wearable-derived SE

| Marker | Model 1<sup>a</sup> |  | Model 2<sup>b</sup> |  | Model 3<sup>c</sup> |  |  |  |
| --- | --- | --- | --- | --- | --- | --- | --- | --- |
|  | Wearable-derived TST |  | Wearable-derived SE |  | Wearable-derived TST |  | Wearable-derived SE |  |
| | $\beta$ | p-value | $\beta$ | p-value | $\beta$ | p-value | $\beta$ | p-value |
| BMI | -0.006 | 0.040 | -0.109 | 0.040 | -0.005 | 0.070 | -0.097 | 0.070 |
| WC | 0.001 | 0.893 | -0.410 | 0.009 | 0.004 | 0.638 | -0.420 | 0.008 |
| WHtR | -3.750 | 0.449 | -0.003 | 0.008 | -2.120x10<sup>-5</sup> | 0.669 | -0.002 | 0.010 |
| RestingHR | -0.014 | 0.004 | -0.024 | 0.800 | -0.015 | 0.004 | 0.011 | 0.910 |
| SBP | -0.009 | 0.493 | -0.180 | 0.461 | -0.008 | 0.550 | -0.161 | 0.513 |
| DBP | -0.008 | 0.383 | -0.011 | 0.952 | -0.008 | 0.384 | 0.009 | 0.960 |
| TotalChol | -0.001 | 0.043 | 0.005 | 0.726 | -0.002 | 0.037 | 0.009 | 0.540 |
| LDL | -0.001 | 0.053 | 0.003 | 0.819 | -0.001 | 0.048 | 0.006 | 0.636 |
| HDL | -9.581x10<sup>-5</sup> | 0.670 | 0.009 | 0.046 | -1.609x10<sup>-4</sup> | 0.517 | 0.010 | 0.039 |
| TG | -2.046x10<sup>-4</sup> | 0.670 | -0.009 | 0.380 | -1.486x10<sup>-4</sup> | 0.779 | -0.008 | 0.404 |
| FBG | -2.354x10<sup>-4</sup> | 0.620 | -0.002 | 0.792 | -2.235x10<sup>-4</sup> | 0.640 | -0.002 | 0.837 |
| BFP | -0.010 | 0.071 | -0.173 | 0.108 | -0.009 | 0.108 | -0.150 | 0.166 |
| SMP | 0.006 | 0.056 | 0.103 | 0.112 | 0.006 | 0.086 | 0.088 | 0.175 |

### Wearable-inferred sleep insufficiency is associated with premature telomere attrition

A subset of the cohort (n = 175) underwent whole-genome sequencing (WGS) for prospective genetic studies. Previous studies have shown that leukocyte telomere length (LTL) can be computationally-inferred from WGS data by analyzing reads containing the telomeric repeat motif (TTAGGG). We used a tool called Telomerecat that estimates LTL by calculating the ratio between read-pairs completely mapping to the telomere and those that span the telomere boundary [35]. WGS-inferred LTL (“WGS-LTL”) was computed for the 175 volunteers with WGS data, and the estimated values were further corrected to account for different sequencing runs. We first confirmed that WGS-LTL values were correlated with volunteer chronological age (r_s_ = -0.264, p = 4.14×10^-04^). When volunteers were split into age-groups of 20-40, 40-60 and 60-80 years, volunteers in the 40-60 and 60-80 years age-groups had WGS-LTL that were on average 259 bp and 427 bp shorter than the reference group of 20-40 years (β = -259.35, p = 0.005 and β = -426.66, p = 0.002 respectively, adjusted for gender and ethnicity, **Fig 3A**). We then asked if wearable-derived sleep metrics were associated with telomere length, and found a positive association between wearable-derived TST and WGS-LTL (β = 1.272, p = 0.020, adjusted for age, gender and ethnicity, **Fig 3B**). We further examined this association by considering two groups of volunteers; those with wearable-derived TST < 5 hours and those with wearable-derived TST > 7 hours. Volunteers with adequate sleep (> 7 hours) had WGS-LTL that was on average 331 bp longer than those with insufficient sleep (< 5 hours) (p = 0.017, adjusted for age, gender and ethnicity, **Fig 3C**). When this analysis was repeated using selfreported TST, a similar association was found between self-reported TST and telomere length (β = 94.404, p = 0.007, **Fig 3D**).

**Fig 3.**
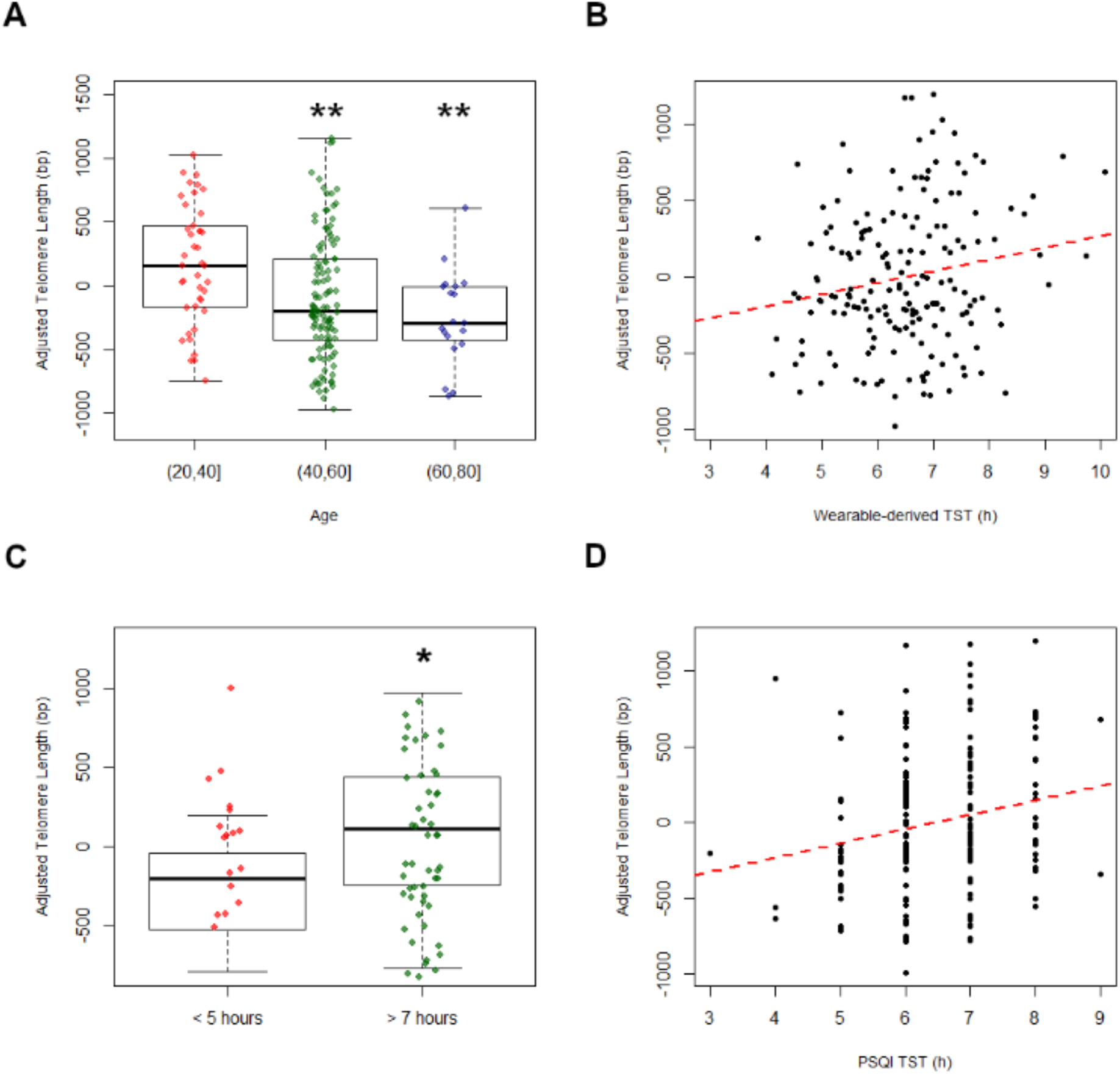
Wearable-derived TST predicts leukocyte telomere length. (A) WGS-LTL by age-group. (B) Wearable-derived TST and WGS-LTL. (C) WGS-LTL of volunteers with insufficient (< 5 hours) and sufficient (> 7 hours) of TST. (D) Self-reported TST and WGS-LTL. The code to generate this figure can be found in S2 Data. Asterisks denote significance of component score in linear model compared to reference score of 0. ** = p < 0.01, *** = p < 0.001. TST = total sleep time; PSQI = Pittsburgh Sleep Quality Index.

## Discussion

We have shown in a sizeable cohort of 482 individuals how sleep metrics obtained from consumer wearables can be used in biomedical research, particularly in the context of health cohort studies. This multi-modal cohort is one of the largest to date to include consumer wearable sleep metrics. Our comparison of objective wearable-derived sleep metrics against subjective measures obtained through the PSQI provides insights into the characteristics of these two modalities. The weak correlation between wearable-derived and self-reported TST is consistent findings from a study that compared between TST obtained from research actigraphs and the PSQI (r = 0.29) [36], despite that study having a considerably longer minimum tracking duration than ours (14 nights vs 3 nights). Concordance was poorer when we compared wearable-derived SE and number of nocturnal awakenings with their selfreported counterparts. As Fitbit devices have been found to perform poorly in terms of detecting wake states [21,24], it is perhaps unsurprising that a significant reduction in wearable-derived SE was only recorded among volunteers reporting the lowest level of SE (< 65%). Our comparison of wearable-derived sleep metrics against volunteer-provided responses to the PSQI – an instrument frequently used in population studies, will inform investigators considering using wearables in future cohort studies. Furthermore, the limitations we highlighted present opportunities for researchers to develop algorithms that can more accurately detect wake states, perhaps by fusing accelerometer and HR data obtained from consumer wearables.

The detailed questionnaires administered as part of our cohort recruitment process afforded us the opportunity to study how wearable-derived sleep metrics are influenced by demographic, socioeconomic and lifestyle factors. Among others, we showed that wearable-derived TST was associated with age, gender, ethnicity, occupation type and even habitual alcohol consumption. That these relationships were not detected previously when self-reported TST was used could be due to several factors. First, wearable-derived TST is quantitative and provides greater resolution compared to self-reported TST, which was in hourly increments. Second, self-reported TST has been shown to be biased and poorly-calibrated compared to objectively-measured habitual sleep duration [37]. With many countries realizing the importance of population health studies, the use of consumer wearables to identify demographic, socioeconomic and lifestyle factors associated with sleep duration could provide vital insights into population sub-groups at risk for poorer health outcomes due to insufficient sleep.

Our analysis on how wearable-derived TST and SE relate to various CVD risk markers demonstrate the utility of wearable-derived sleep metrics in biomedical research. Xu et al. previously described a link between habitual sleep duration estimated using Fitbit Charge HR devices and BMI in a predominantly European-American cohort of 471 individuals. We identified in an Asian cohort this relationship despite a considerably shorter average tracking duration (4 nights vs 78 nights), suggesting that sleep metrics obtained from even short studies can be useful. One novel aspect of this study was our analysis of wearable-derived SE against CVD risk markers. Our findings of links between SE and three obesity markers; BMI, WC and WHtR, is supported by previous studies performed using orthogonal approaches such as PSG and actigraphy [38,39]. This indicates that wearables are capable of contributing beyond TST to health cohort studies, notwithstanding current limitations to the accuracy of wearable-derived SE. The paucity of associations between wearable-derived sleep metrics and clinical parameters such as fasting blood glucose (a marker of insulin sensitivity), and blood pressure could be due in part to the cohort size, and highlights the need for wearables to be included in larger population-scale cohort studies in order to thoroughly assess their utility.

Beyond comparisons with the usual clinical health markers, our study provides a novel demonstration of the utility of wearable-derived sleep data in the study of biological aging. In particular, we showed that in a normal free-living cohort, individuals with short habitual sleep duration also exhibited premature telomere attrition. Previous studies of this phenomenon have either used sleep questionnaires [31–34] or cohorts of sleep disorder patients [40,41]. As premature telomere shortening has been linked to the early onset of various age-related diseases [42] and all-cause mortality [43,44], new evidence on the link between insufficient sleep and accelerated aging such as this are vital in helping shape public policy (e.g. later school start times, altered work hours and schedules, etc.) and to promote healthier sleep habits among the public. This finding is especially relevant to Singapore, a developed nation whose citizenry is among the most sleep-deprived in the world [45,46]. Our use of 1) wearables for sleep tracking and 2) WGS for LTL estimation in this study, demonstrate the versatility that emerging technologies can bring to population cohort studies by providing added behavioral and phenotypic data beyond their primary functions.

In summary, our study has demonstrated various aspects in which sleep metrics from wearables can be used in cohort studies. Apart from comparing wearable-derived and selfreported sleep metrics, our work has shown that wearables can be used to study how sleep relates to demographic, socioeconomic and lifestyle factors, as well as various markers of health and aging. The increasing ubiquity of wearables and other forms of digital health, represent a rich source of behavioral data that can be tapped by investigators running cohort studies. Beyond the use of wearables as study-provided devices, a BYOD (bring your own device) model, where participants share data from their own wearables with investigators through application programming interfaces (APIs), is also possible. This is particularly attractive as the BYOD model allows for much longer tracking durations with minimal incremental cost. At the same time, the use of wearables and other digital health devices in population health studies can catalyze further development of digital applications that promote healthy behavior, including sleep habit.

## Materials and Methods

### Study volunteers and ethics statement

Volunteers were recruited as part of the SingHEART/Biobank study using a protocol and written informed consent form approved by the SingHealth Centralized Institutional Review Board (ref: 2015/2601). Details on the cohort and its inclusion criteria is available in [26]. Among others, the cohort underwent an activity tracking study a consumer wearable (Fitbit Charge HR), detailed clinical profiling for various CVD risk markers including anthropometry, blood pressure, lipid panel and fasting blood glucose. After evaluation for completeness of sleep and activity tracking data, and the removal of subjects with extreme outlier activity metrics, 482 volunteers were included in this study.

### Processing of wearable sleep metrics

For each volunteer, we extracted their Fitbit data (activity, heart rate and sleep) using the Fitbit Web API (https://dev.fitbit.com/reference/web-api/quickstart/). Data completeness was evaluated by the availability of HR data and days with no intraday steps were excluded (full details are available in [26]). We considered days with at least 20 hours of data to be complete, and only volunteers with at least three data-complete days were included. Detailed sleep tracking data from Fitbit was obtained in the JSON (JavaScript Object Notation) format, and processed using an R script (available in **S2 Data**). For each day, we summed the duration of all sleep sessions starting between 8 PM and 8AM. We then averaged daily sums for each volunteer to obtain the TST. Sleep hour was determined by calculating the average start time of sleep sessions occurring between 8 PM and 8 AM with duration more or equal to 3 hours. Wake hour was determined by averaging the end time of sleep sessions. SE was computed in a similar fashion, except that for each day, the average SE of sleep sessions was obtained. In addition, we estimated the number of nocturnal awakenings by averaging daily total wake counts.

### Questionnaires

The volunteer recruitment process included the administration of several questionnaires. This included the SingHEART patient questionnaire which covered demographics, socioeconomic factors, medical history, smoking history, alcohol consumption patterns, exercise and dietary habits. The General Practice Physical Activity Questionnaire (GPPAQ) was used to estimate physical activity levels (details on how we converted GPPAQ responses to numerical scores are available in [26]). Finally, the Pittsburgh Sleep Quality Index (PSQI) questionnaire, which assesses sleep quality and disturbances over a one-month time interval, was also administered [29]. Volunteers were asked about their sleep habits, for example their bed time, hours of sleep per night, sleep trouble and sleep quality. The PSQI contains 19 self-rated questions and 5 questions rated by the bed partner or roommate (if one is available). These 19 self-rated items are then combined to produce seven component scores, each of which has a range of 0-3. Components 3 and 4, as well as Question 5b were examined in this study due to their relevance to our wearable-derived sleep metrics (TST, SE and nocturnal awakenings respectively).

### Association tests

Multiple linear regression analyses described in this study were conducted using the GLM (generalized linear model) function in R and Gaussian error distribution was used. When gender was considered as a covariate, the female gender was set as the reference level. Whereas the following are the reference level for each of the socioeconomic factor when it was set as a covariate: “Public-housing” for residence type; “Others” for education level and “Manual-labor” for occupation type. For lifestyle factors, the reference level for alcohol and caffeine, tea and green tea consumption is “No”; the reference levels for smoking and exercise status are “Ex-smoker” and “Never/hardly” respectively.

For linear regression analysis between wearable-derived sleep metrics and CVD risk markers, three models were used. Age, gender, ethnicity, daily step counts were included as covariates. For Model 1 and Model 2 which considered TST and SE-based sleep metric, the model is, respectively, Marker ~ Age + Gender + Ethnicity + AverageDailyTotalSteps + TST and Marker ~ Age + Gender + Ethnicity + AverageDailyTotalSteps + SE. Whereas for Model 3 which includes both TST and SE, the model is Marker ~ Age + Gender + Ethnicity + AverageDailyTotalSteps + TST + SE. For linear regression analysis between wearable-derived TST metrics and socioeconomic factors, the following model was used: TST ~ Age + Gender + Ethnicity + SocioeconomicFactor. In addition, linear regression analysis between wearable-derived TST metrics and LTL was done by using this model: LTL ~ Age + Gender + Ethnicity + TST.

### Telomere length estimation

Telomerecat [35] was used to estimate LTL from WGS data (WGS-LTL) by calculating the ratio between read-pairs completely mapping to the telomere and those that span the telomere boundary. As part of a larger study, we had sequenced the genomes of 546 volunteers (of which 175 overlapped with this study) at a target depth of 30X. We used Telomerecat to estimate WGS-LTL for the volunteers. Briefly raw sequencing reads in FASTQ files were aligned to the human reference genome (hs37d5) using BWA-MEM version 0.7.12 [47]. The resulting BAM files were further processed using Sambamba version 0.5.8 [48] to sort the reads and flag duplicates. Telomerecat was run in two steps. First, the *bam2telbam* command was run on individual BAM files to generate *telbam* files, which are small BAM files containing only sequencing reads relevant to LTL estimation. The *telbam2length* command was then executed to generate LTL estimates for the entire 546 set of *telbam* files. Finally, we used linear regression to correct for different sequencing runs. BAM files containing telomeric and subtelomeric sequencing reads are available in the European Nucleotide Archive (ENA, accession number PRJEB29577).

### Software and statistical tests

All statistical analyses in this study were performed using the R statistical environment. Unless otherwise stated, correlations described in this study are Spearman correlation coefficients.

## Data and code availability

All data (including raw wearable metrics) and R code used in this study are available in S2 Data.

## Acknowledgments

We would like thank the volunteers for their participation, as well as the NHCS and SingHEART Clinical Research Coordinators.

## Author contributions

J.X.T., K.K.Y., P.T. and W.K.L. conceived the study. W.K.L, K.K.Y. and P.T. directed the study and supervised the research. J.X.T., W.K.L., S.D. and S.G.R. performed bioinformatics and biostatistical analysis. C.Y. and C.J.P. were involved in sample preparation and processing. J.Y., S.Y.T., S.A., C.W.C. and B.T.T. performed interpretation of clinical data and findings. J.X.T and W.K.L., K.K.Y. and P.T. wrote the manuscript, with the assistance and final approval of all authors.

## Supporting Information

**Supplementary Table 1.** Summary of multiple linear regression results for socioeconomic factors and sleep metric, wearable-derived TST. β-values, P-values and standard error are shown. Highlighted cells have p < 0.05. The reference level in each model is stated as “ref” and highlighted in grey.

**Supplementary Table 2.** S2 Table: Summary of multiple linear regression results for lifestyle factors and sleep metric, wearable-derived TST. β-values, P-values and standard error are shown. Highlighted cells have p < 0.05. The reference level in each model is stated as “ref” and highlighted in grey.

**S1 Data.** Characteristics of study participants.

**S2 Data.** All data and R code used in this work.

